# DeepAction: A MATLAB toolbox for automated classification of animal behavior in video

**DOI:** 10.1101/2022.06.20.496909

**Authors:** Carl Harris, Kelly R. Finn, Peter U. Tse

## Abstract

The identification of behavior in video is a critical but time-consuming component in many areas of animal behavior research. Here, we introduce DeepAction, a deep learning-based toolbox for automatically annotating animal behavior in video. Our approach uses features extracted from raw video frames by a pretrained convolutional neural network to train a recurrent neural network classifier. We evaluate the classifier on two benchmark rodent datasets and show that it achieves high accuracy, requires little training data, and surpasses both human agreement and similar existing methods. We also create a confidence score for classifier output, and show our method provides an accurate estimate of classifier performance and reduces the time required by human annotators to review and correct automatically-produced annotations. We release our system and accompanying annotation interface as an adaptable, non-technical, and open-source MATLAB toolbox.

## Introduction

The classification and analysis of animal behavior in video is a ubiquitous but often laborious process in life sciences research. Traditionally, such analyses have been performed manually. This approach, however, suffers from several limitations. Most obvious is that it requires researchers to allocate much of their time to the tedious work of behavioral annotation, limiting or slowing the progress of downstream analyses. Particularly for labs without research assistants or paid annotators, the opportunity cost of annotating video can be quite high. Manual annotation also suffers from relatively poor reproducibility and reliability^1–3^, largely due to the limited attention capacity of human annotators. This issue is particularly salient in studies involving rodents. Due to their nocturnal nature, rodents are preferably studied under dimmed or infrared light^4^, which makes the identification of behaviors more difficult due to more limited light and color cues. This, in turn, increases annotators’ fatigue and reduces their capacity to pay attention for extended periods, introducing variation in annotation quality across time and space conditions that decreases the quality of behavior data^5^.

Given the time and accuracy limitations of manual annotation, increasing work has focused on creating methods to automate the annotation process. Many such methods rely on tracking the animals’ bodies^4,6–9^ or body parts^10^, from which higher-level features (e.g., velocity, acceleration, and posture) are extrapolated and used to classify behavior. Jhuang, et al. ^7^, for example, used motion and trajectory features to train a hidden Markov support vector machine to categorize eight classes of mouse behavior. Burgos-Artizzu, et al. ^6^ used spatiotemporal and trajectory features and a temporal context model to classify the social behavior of mice using two camera views. However, approaches using these “hand-crafted features” are limited in several ways^11^. First, they require researchers identify sets of features that both encompass a given animal’s entire behavioral repertoire and can distinguish between visually similar behaviors. For example, “eating” and “grooming snout” behaviors in rodents do not have a well-defined difference in posture or movement^4^, making crafting features to differentiate them difficult. Second, after features have been selected, detecting and tracking them is difficult and imperfect. Subtle change in video illumination, animal movement, and environment can result in inaccurate keypoint detection, decreasing the fidelity of extracted features. And third, selected feature sets are often experiment-specific. Those optimal for a singly housed rodent study, for example, likely differ from those optimal for a social rodent study. This increases the complexity of the feature-selection task, impeding experiment progress and annotation accuracy.

To address these limitations, Bohnslav, et al. ^11^ proposed an alternative to hand-crafted approaches, instead using hidden two-stream networks^12^ and temporal gaussian mixture networks^13^, and achieved high classification accuracy on a diverse collection of animal behavior datasets. Here, we expand on this work by introducing DeepAction, a MATLAB toolbox for the automated annotation of animal behavior in video. Our approach utilizes a two-stream^14^ convolutional and recurrent neural network architecture^15,16^ to generate behavioral labels from raw video frames. We use convolutional neural networks (CNNs) and dense optical flow to extract spatial and temporal features from video^17^, which are then used to train a long short-term memory network classifier to predict behavior. We evaluate our approach on two benchmark datasets of laboratory mouse video and show it outperforms existing methods and reaches human-level performance with little training data. In addition to outputting behavior labels for each video frame, we also introduce a classification confidence system that generates a measure of how “confident” the classifier is about each label. This allows researchers to estimate the quality of automatically-produced annotations without having to review them, and reduces the time required to review annotations by allowing users to selectively correct ambiguous ones, while omitting those the classifier produced with high confidence. We show this confidence score accurately differentiates low quality annotations from high quality ones and improves the efficiency of reviewing and correcting video. Finally, we release the code and annotation GUI as an open source, accessible, and adaptable toolbox.

## Results

### The DeepAction workflow

The toolbox workflow (**Fig. 1A**) begins with the importation of unlabeled video into a new DeepAction project and ends with the export of annotations for all the videos in that project. The workflow consists of two parts: a classification component (steps 2-8) and a review component (steps 9 and 10). In the classification portion, we adopt a supervised learning approach in which a portion of project videos are labeled and used to train a classifier. This classifier learns to associate the content in the video frames with a set of user-defined behavior labels (e.g., “walk” or “drink”). After the classifier is trained, it can then be used to predict behaviors in the *unlabeled* video. In addition to predicting behaviors occurring in the unlabeled video, the classifier outputs a “confidence score,” representing an estimate of the agreement between classifier-produced labels and human-produced ones. This confidence score is used during review component of the workflow, in which low-confidence annotations can be preferentially reviewed and corrected, while those with high confidence are omitted. After this confidence-based review, annotations are exported for use in the researcher’s given analysis.

**Figure 1:**
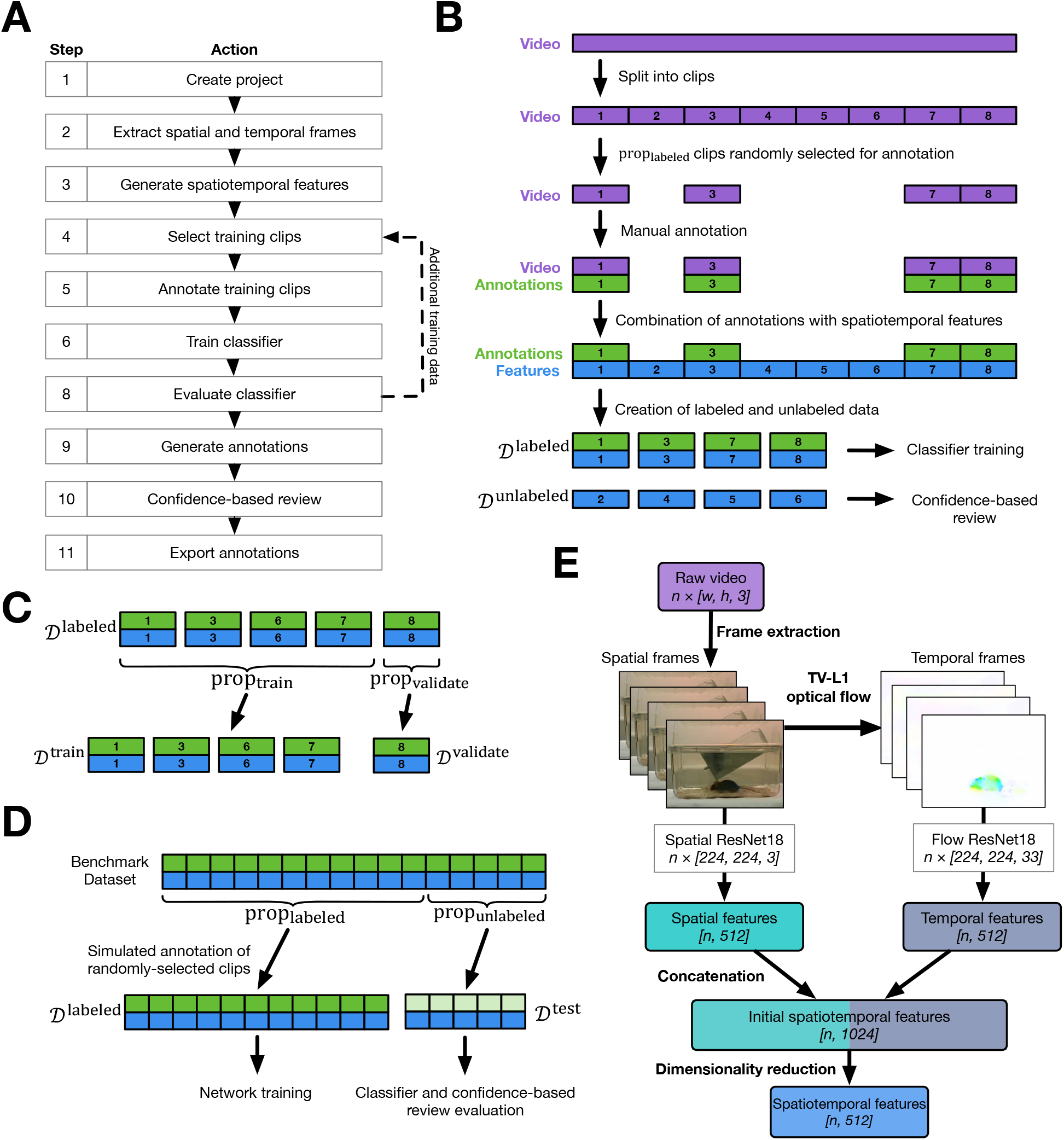
Toolbox workflow and data selection process. (**A**) Workflow for the DeepAction toolbox. Arrows indicate the flow of project actions, with the dashed arrow denoting that, following training the classifier, additional training data can be annotated and used to re-train the classifier. (**B**) An overview of the clip selection process. Long videos are divided into clips of a user-specified length, from which a user-specified proportion (prop_labeled_) are randomly selected for annotation 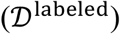. The selected video clips are then annotated, and these annotations are used in combination with their corresponding features to train the classifier. The trained classifier is used to generate predictions and confidence scores for the non-selected clips 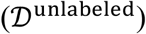, which the user can then review and correct as necessary. (**C**) Labeled data are further divided into training 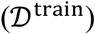 and validation 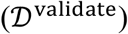 data. (**D**) Process for simulating clip-selection using our benchmark datasets, where we simulate selecting prop_labeled_of the data for labeling 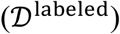 and evaluate it on the unselected data 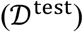. (**E**) Process to generate spatiotemporal features from video frames. Raw video frames are extracted from the video file (“frame extraction”). The movement between frames is calculated using TV-L1 optical flow and then represented visually as the temporal frames. Spatial and temporal frames are input into their corresponding pretrained CNN (“spatial ResNet18” and “flow Resnet18,” respectively), from which spatial and temporal features are extracted. The spatial and temporal features are then concatenated, and then their dimensionality is reduced to generate the final spatiotemporal features that are used to train the classifier. Dimensionality of is shown in italicized brackets.

To represent the video for input to the classifier, we opt for a “two-steam” model^18^, where the first stream (“spatial stream”) captures the spatial information of the video frames, and the second stream (“temporal stream”) captures the motion between frames (**Fig. 1E**). We first extract video frames representing the spatial and temporal information (“spatial frames” and “temporal frames,” respectively) in the underlying video (see **Methods: Frame Extraction**). To generate spatial frames, which contain information about the scenes and objects in the video, we extract the raw video frames from each video file. To generate temporal frames, which contain information about the movement of the camera and objects in the video, we use dense optical flow to calculate the movement of individual pixels between *pairs* of sequential frames. Dense optical flow generates a two-dimensional vector field for each pixel in the image, where each vector represents the estimated movement of a pixel from one image to the next^19^. We then express this entire vector field visually as an image, where a given pixel’s color is governed by the orientation and magnitude of its corresponding flow vector.

We then generate a low-dimensional representation of the spatial and temporal frames by extracting their salient visual features using the ResNet18 pretrained convolutional neural network (CNN; see **Methods: Feature extraction**). For each spatial and temporal frame, the feature extractors generate a 512-dimensional vector representing the high-level visual information contained in that frame. We then concatenate these spatial and temporal features (dimensionality of 1024) to create the initial spatiotemporal features, and then use reconstruction independent component analysis to reduce the dimensionality to 512, forming the final spatiotemporal features used to train the classifier.

Training the classifier requires a portion of video be manually labeled so it can learn the associations between the video’s corresponding spatiotemporal features (input) and behavior labels (output). Rather than annotating whole videos at a time, we instead split each video into short “clips,” where each clip is a short segment of the longer video, and then select a subset of these clips to annotate (**Fig. 1B**). This approach is preferable because it better captures the substantial variation in features and the feature-to-label relationship *across* videos, improving the generalizability of the classifier. As long as clips are long enough to annotate accurately, the user experience is similar, provided total annotation time remains the same.

After the set of videos has been split into clips, a portion of these clips specified by the user, prop_validate_, are randomly selected for manual annotation (**Fig. 1B**) using a GUI included in the toolbox release (**Fig. 6B**). After annotation, labeled clip data (video, features, and annotations), 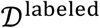, is used to train a recurrent neural network classifier (see **Methods: Classifier architecture**) and the confidence-based review system. To do so, we first further split 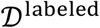 into a training set and a validation set (**Fig. 1C**). The training set, 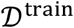, comprises most of the labeled data and is directly used to train the classifier. For a given clip in 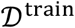 with *n* frames, a spatiotemporal feature array of size [*n,* 512] is input into a recurrent neural network classifier, along with a series of *n* manually annotated behavior labels. The network then tries to predict the manual annotations using the features; training iteratively reduces the difference between classifier-predicted and human annotations. The spatiotemporal features corresponding for a given segment of video represent the visual content of that segment; so, by predicting labels using these features, the classifier is indirectly generating predictions for the underlying video data. The independent validation set, 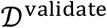, is used to tune the model training process and confidence-based review (see **Methods: Classifier training**). The trained classifier and confidence-based review system are then used to generate annotations and confidence scores for the remaining, unlabeled data, 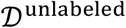.

We then introduce a confidence-based review system. Recall that, after the classifier has been trained, it can be used to predict behaviors in unlabeled data, 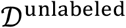. In addition, we output a confidence score for each clip in 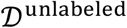 corresponding the estimated accuracy of the labels produced for that clip (see **Methods: Clip confidence definition**). In an ideal metric, a clip’s confidence score should correspond to the ground truth likelihood of the classifier-predicted behaviors being correct. The purpose of the confidence score is two-fold (see **Methods: Confidence-based review**). First, by generating estimated accuracies for each clip in 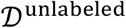, we can estimate the *overall* accuracy of 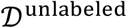. Just as there is variability in annotations between researchers, we can expect that even a well-performing classifier’s annotations will not exactly match those that would be produced if the unlabeled data was manually annotated. But, by providing an estimate of the agreement between human- and classifier-produced labels in 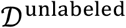 automatically, users can easily decide whether the classifier’s performance is sufficient for their given application. The second purpose is to enable researchers to preferentially review and correct clips where the classifier is less accurate over those where annotations are highly accurate. Rather than reviewing each clip in 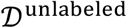, researchers can review and correct only the subset of the clips where the classifier is uncertain about its predictions. If the confidence score is a precise estimate of accuracy, then the clips with a low confidence score will be the clips that the classifier performs poorly on, allowing for labels to be corrected more efficiently.

### Datasets

We evaluate our approach on two publicly available datasets of mice in a laboratory setting (see **Methods: Datasets**). Both datasets are fully annotated, allowing us to test and evaluate our model. The first dataset, referred to as the “home-cage dataset,” was collected by Jhuang, et al. ^7^, and features 12 videos (10.5 hours total; **Fig. 2D**) of singly housed mice in their home cages performing eight stereotypical, mutually-exclusive behaviors recorded from the side of the cage (**Fig. S1A**). The second dataset, called “CRIM13,”^6^ consists of 237 pairs of videos, recorded with synchronized side and top views, of pairs of mice engaging in social behavior, categorized into 13 distinct, mutually-exclusive actions (**Fig. S1B**). Each video is approximately 10 minutes each, for a total of approximately 88 hours of video and annotations (**Fig. 2D**).

**Figure 2.**
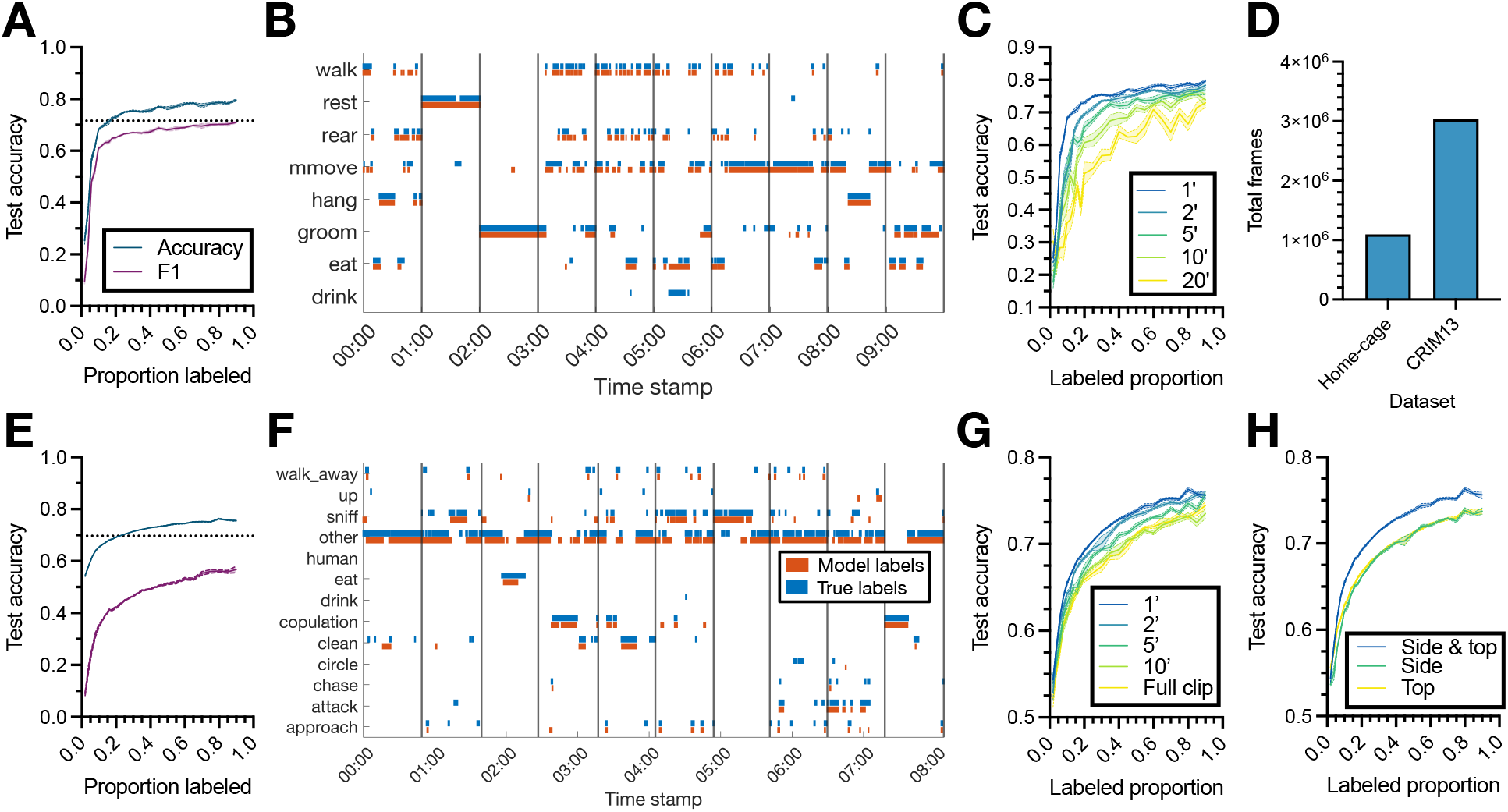
Classifier performance. (**A**) Test set accuracy and overall F1 score of the classifier on the home-cage dataset as a function of the proportion of the dataset used to train it. The proportion of data denoted on the *x*-axis is used to train the classifier, which is then evaluated on the remainder of the dataset. (**B**) Sample ethogram of classifier labels and ground-truth annotations from 10 randomly selected home-cage clips. Each colored line indicates the label of that behavior at the corresponding time stamp. Vertical black lines denote the divisions of the video into clips (one minutes in duration). (**C**) Test set accuracy on the home-cage dataset as a function of the proportion of data used to train the classifier, for clips of varying length (clip duration denoted in minutes). (**D**) Total number of annotated frames in each dataset. (**E-G**) Same as (**A-C**),but for the CRIM13 dataset. (**H**) Test set accuracy on the CRIM13 dataset as a function of the amount of training data for classifiers trained with features from the side camera, top camera, and both the side and top cameras. Lines and shaded regions in (**A,C,E,G,H**) indicate mean and standard error, respectively, across 10 random splits of the data.

Because both datasets are already fully labeled, we evaluate our method by simulating the labeling process (see **Methods: Simulating labeled data**). We assume a user has chosen to annotate some proportion of the data and uses that data to train a classifier and obtain predictions for the remaining data. In practice, the user would run the classifier over the remaining data to automatically generate labels and use the confidence-based review system to review those labels as desired. Here, however, since the data have been annotated, we know the true labels for the “unlabeled” data. This allows us to test the performance of the method *as if it were being used* to produce labels for the remaining prop_unlabeled_ data. This approach allows us to simulate performance across a range of labeling proportions, which in turn provides a measure of how to model can be expected to perform for a given amount of manual annotation time. So, for a given prop_labeled_, we train the classifier and confidence-based review using the “labeled data,” and then test how the approach performs on the remaining data (**Fig. 1D**).

### High classification accuracy with little training data

We first evaluate the performance of the classifier (i.e., accuracy and F1; see **Methods: Classifier evaluation**) with varying amounts of training data (**Fig. 2A,E**), and show that it requires remarkably little manual annotation to achieve high accuracy. For a given proportion labeled (i.e., prop_unlabeled_ above), a corresponding proportion of project clips are randomly selected from all the clips in the dataset and used to train the classifier, which is then evaluated on the remaining data (i.e., the test set). For both datasets, accuracy and F1 improve as more training data is used, with steep increases for the first ten percent of the data, and more gradual increases after twenty percent. Example classifier output and ground truth annotations are shown in **Figure 2B,F**.

We then compare the performance of our model with existing ones (**Table 1**; see **Methods: Comparison with existing methods**). On the home-cage dataset, in addition to showing higher accuracy than the agreement between human annotators, our classifier outperforms existing commercial options, as well as approaches based on hand-crafted features^7^ and 3D convolutional neural networks^20^. The classifier also demonstrates above-human performance and surpasses the sparse spatio-temporal feature approach detailed in Burgos-Artizzu, et al. ^6^ on the CRIM13 dataset.

**Table 1.** Performance comparison with existing methods. Shown is the accuracy of various annotation methods on both datasets. “Human” denotes the agreement between two human annotator groups (see **Methods: Inter-observer reliability**). The accuracy for DeepAction on the home-cage and CRIM13 datasets is the mean accuracy from 12-fold and two-fold cross-validation, respectively, to provide a comparable reference to Jhuang, et al. ^7^ and Burgos-Artizzu, et al. ^6^ (see **Methods: Comparison with existing methods**).

|  | Model | Accuracy |
| --- | --- | --- |
| Home-cage | Human | 71.6% |
|  | CleverSys commercial system<sup>7</sup> | 61.0% |
|  | Jhuang, et al.<sup>7</sup> | 78.3% |
|  | Murari<sup>20</sup> | 73.5% |
|  | **DeepAction** | **79.5%** |
| CRIM13 | Human | 69.7% |
|  | Burgos-Artizzu, et al.<sup>6</sup> | 62.6% |
|  | **DeepAction** | **73.9%** |

### Data input process improves performance

Next, we consider how unique aspects of our data preparation process affect the performance of the classifier. Specifically, we investigate our hypothesis that, given equal annotation time (i.e., an equal labeled proportion), our classifier shows superior performance when it is trained using relatively short clips rather than longer ones. As shown in **Figure 2C,G**, this is indeed the case. While presumably there is a limit to this phenomenon (i.e., if the clip length were to be only a handful of frames, the classifier would fail to gain enough context to accurate predict its labels), in the clip durations tested here, varying between one and 20 minutes, shorter clips are both more accurate for a given level of annotation and demonstrate a more rapid improvement as training data increases. The CRIM13 dataset is recorded using synchronized top- and side-view cameras. In our main analysis we combine the features from both cameras (see **Methods: Feature extraction**); in **Figure 2H** we confirm that this is advantageous. The classifier trained using features from both views demonstrates superior performance to one trained only features from the side camera or only those from the top camera, indicating our method effectively integrates information from multiple cameras.

### DeepAction performs well across behaviors

An important consideration, in addition to overall classifier performance, is classifier performance on specific behaviors. In highly imbalanced datasets (i.e., those in which a small number of behaviors are disproportionally common), high accuracy can be achieved by a classifier with poor discriminative capacity if its predictions are the most common classes. The home-cage dataset, except for the “drink” behavior (0.26 percent of labels), is relatively well-balanced (**Fig. 3A**). For non-drinking behaviors, the classifier shows consistently high performance (**Fig. 3B**), despite modest variation in the prevalence of each label. The CRIM13 dataset displays significantly less balance (**Fig. 3D**), with a high proportion of behaviors classified as “other” (denoting non-social behavior). The high incidence of the “other” behavior accounts for the high performance of the classifier at near-zero training data proportions (approximately 55 percent accuracy; **Fig. 2E**), and a disproportionately large number of social behaviors being incorrectly labeled as “other” by the classifier (**Fig. 3F**). We also note that the distribution of bout lengths (i.e., the number of frames for which a behavior consecutively occurs) predicted by the classifier is qualitatively similar to the true distribution of bout length for most behaviors. In the home-cage dataset we see that the classifier underpredicts bout lengths for the “rest” behavior, which has an exceptionally long average bout length (2,563 frames vs. an average of 88 frames for all other behaviors), despite its high performance in predicting the rest behavior overall (recall: 0.95, precision: 0.98) on the same test set. In the CRIM13 dataset, we observe that the classifier underpredicts bout lengths for the behaviors it performs worst on: “eat,”“human,” and “drink.”

**Figure 3:**
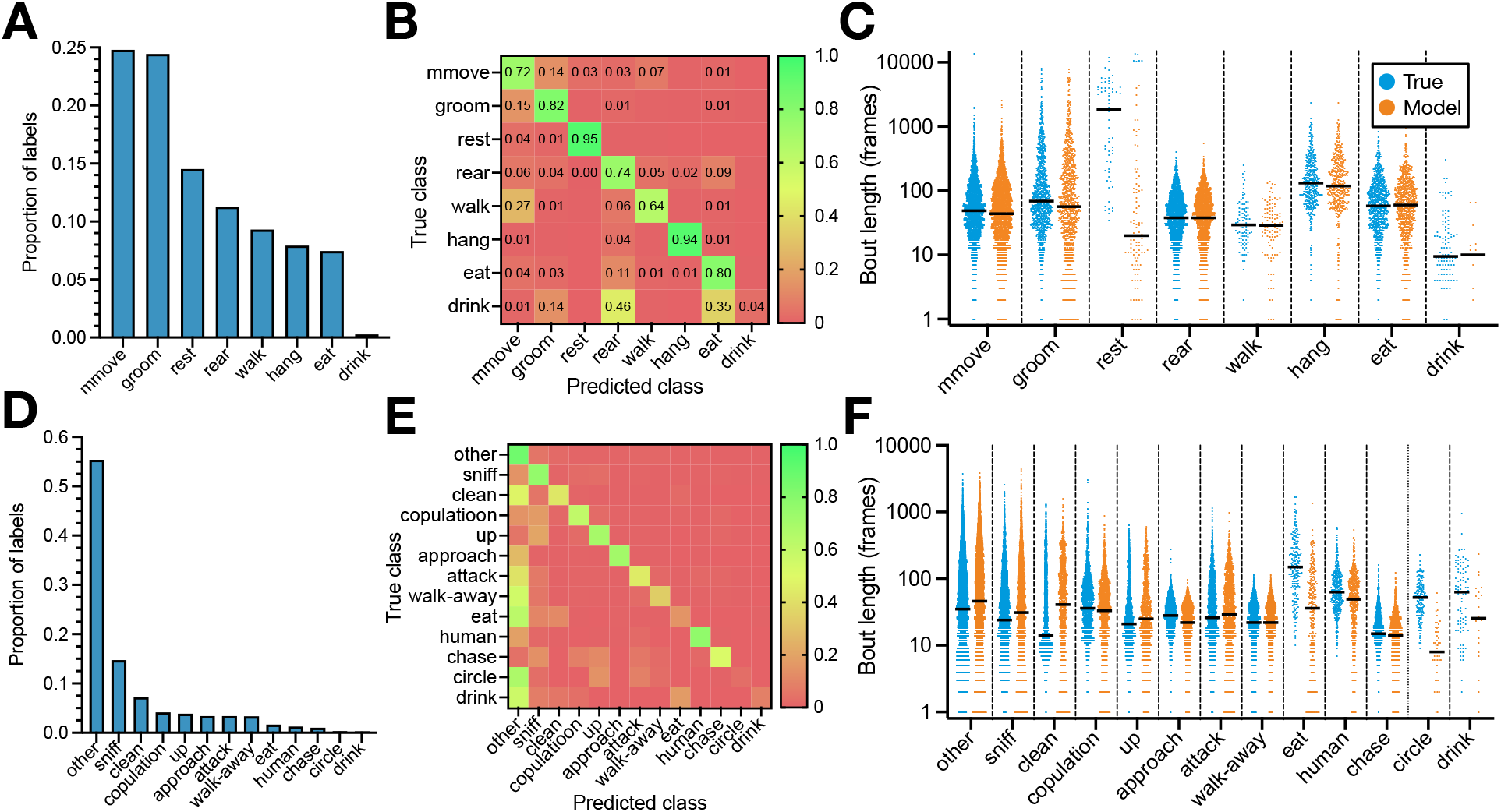
Dataset behavior characteristics and classifier performance. (**A**) Ground-truth distribution of behavior labels (i.e., the number of frames in which each behavior occurs as a proportion of the total number of frames in the dataset) for the home-cage dataset. (**B**) Example confusion matrix showing the classifier performance by behavior on the home-cage dataset, with cell values normalized relative to the true class. (**C**) True bout lengths and example predicted bout lengths for the home-cage dataset, grouped by behavior. A single “bout” refers to a period of continuously occurring behavior, and the corresponding bout length to the length of that period in number of frames. Median bout length is marked by the solid black lines, and each dot corresponds to a single bout. (**D-F**) Similar to (**A-C**), but for the CRIM13 dataset.

To examine classifier performance as a function of the amount of data used to train it, we calculate the precision, recall, and F1 score (see **Methods: Classifier evaluation**) for each behavior with varying labeled data proportions (**Fig. 4**). In the home-cage dataset, for non-drinking behaviors, we observe a similar pattern in behavior-level improvement as we do to overall accuracy - a rapid increase at low training data proportions, followed by a more gradual one at 10 to 20 percent training data (**Fig. 4A-G**). This pattern holds even given the relatively large difference in incidence between the least common (eat, at 7.5 percent of labels) and most common (micromovement, 24.8 percent of labels) non-drink behaviors. For drinking behavior, however, due to its exceptionally low incidence, we observe a more inconsistent, non-gradual improvement in performance across training set proportions (**Fig. 4H**).

**Figure 4.**
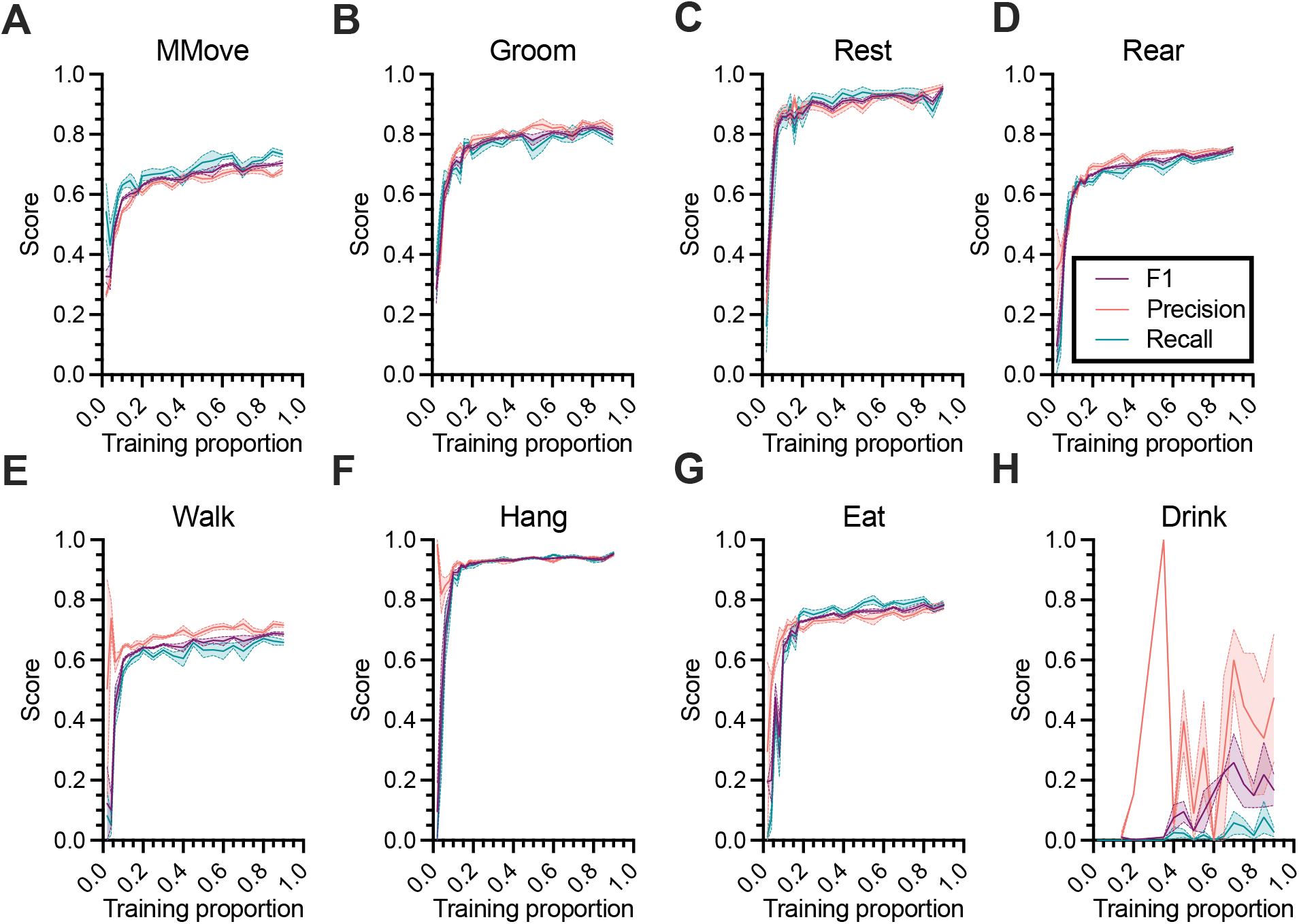
Home-cage behavior-level classifier performance. (**A-H**) Precision, recall, and F1 scores for each behavior in the home-cage dataset as a function of the proportion of data used to train the classifier. Lines and shaded regions indicate mean and standard error, respectively, across 10 random splits of the data.

This pattern generally applies in the CRIM13 dataset as well (**Fig. S3**). For most behaviors we observe a rapid increase in recall, precision, and F1, followed by a relative slowdown in improvement as a function of training proportion at a training proportion of approximately 0.3. There are notable exceptions to this pattern. First, we observe that, as compared to very low training proportions, the recall of “other” decreases slightly as the classifier defaulted to predicting “other” with disproportionate frequency (**Fig. S3A**). The F1 score, however, increased, indicating and improved balance between recall and accuracy. And second, we observe that “eat,” “circle,” and “drink” show sporadic improvements in recall, precision, and F1 as a function of training proportion (**Fig. S3I,L,M**). As with “drink” in the home-cage dataset, these are all low-incidence behaviors (approximately 2 percent of ground-truth labels or less), particularly in the case of “circle” and “drink” (approximately 0.3 percent of ground-truth labels).

### Calibrated confidence scores accurately predict classification accuracy

Next, we turn our focus from the performance of the classifier to the performance of the confidence-based review. Recall that we generate a confidence score for each clip that represents the classifier’s prediction of the accuracy of its predicted labels (see **Methods: Confidence score definition**). In **Figure 5A,D** we demonstrate that there is a strong correlation between confidence score and accuracy, for both confidence scores based on maximum softmax probability and those derived using temperature scaling (see **Methods: Confidence score calculation**). We next consider the mean absolute error (MAE; see **Methods: Evaluating confidence score calibration**) between clips’ predicted accuracy (i.e., confidence score) and actual accuracy across training data proportions. Here, the MAE expresses the amount by which a randomly selected clip’s confidence score differs (whether positively or negatively) from its accuracy. The MAE derived using temperature scaling performs slightly better than the one derived using softmax probabilities on the CRIM13 dataset (**Fig. 5E**) but not the home-cage dataset (**Fig. 5B**). While the MAE for both methods improves initially, it plateaus after the proportion of data labeled reaches about 20 percent, indicating that exact estimates of clip accuracy remain elusive.

**Figure 5.**
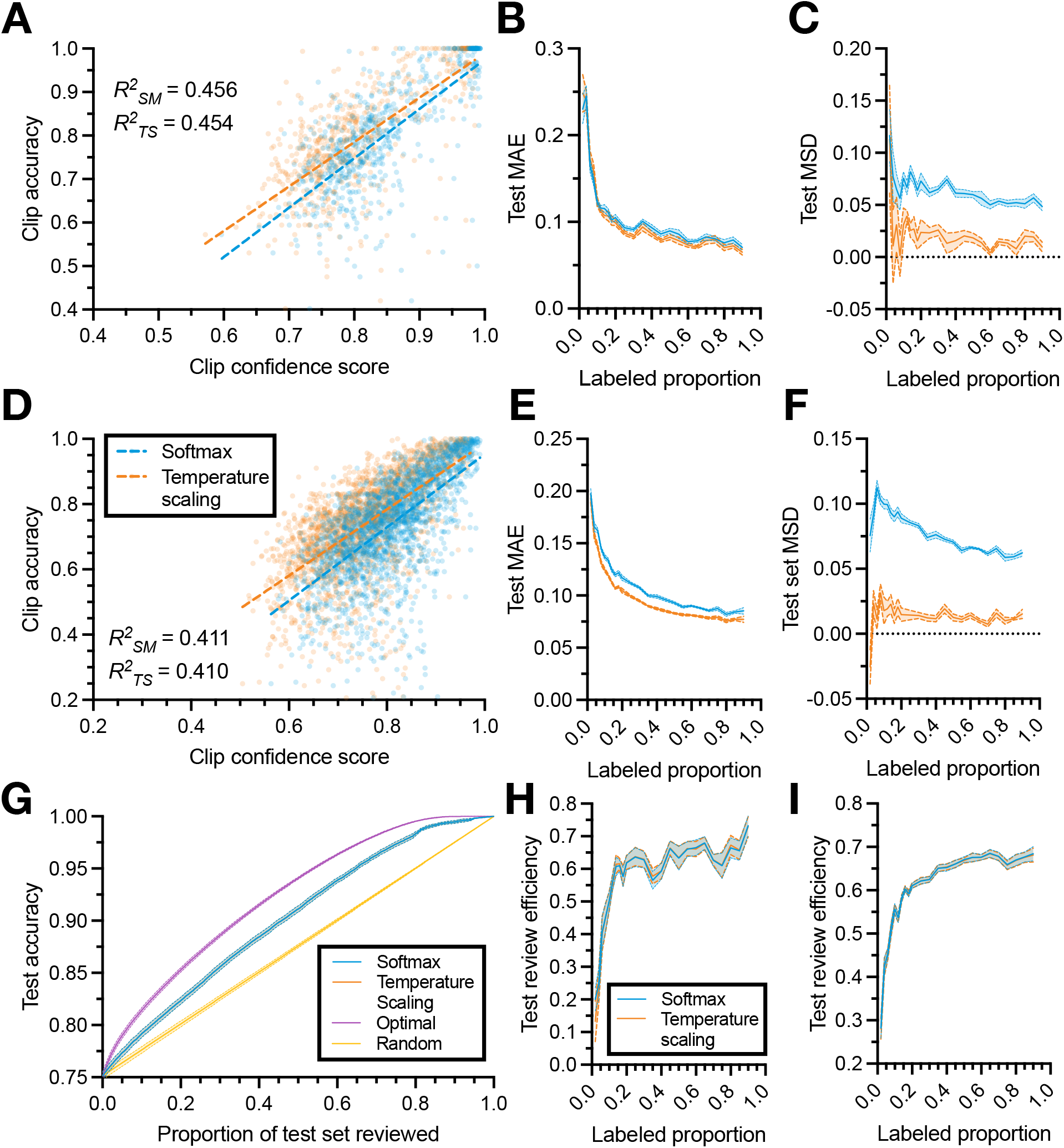
Confidence measure improvements across training proportions. (**A**) Example of the correlation between clip confidence score and clip accuracy. Dashed lines indicating the line of best-fit with r-squared values inset. (**B**) Mean absolute error (MAE) and (**C**) mean signed difference (MSD) between clip confidence score and clip accuracy as a function of the amount of data used to train the classifier. (**D-F**) Similar to (**A-C**), but for the CRIM13 dataset. (**G**) Example relationship between the proportion of test clips reviewed (and corrected) and test set accuracy from the home-cage dataset, where clips are reviewed in an order determined by the confidence scoring method, for various scoring methods (see **Methods: Confidence-based review**). (**H**) Review efficiency metric, quantifying how effectively a given confidence scoring method refers low-confidence clips for review first (see **Methods: Evaluating review efficiency**) as a function of the amount of training data, for the home-cage dataset. (**I**) Same as (**H**), but for CRIM13. Lines and shaded regions in (**B,C,E,F,H,I**) indicate mean and standard error across 10 random splits of the data.

Perhaps more important than predicting the accuracy of classifications on a single clip is predicting the accuracy of classifications across all unlabeled clips. While the absolute error of individual clips might fluctuate, if the differences cancel out (i.e., if predictions are just as likely to be overconfident as they are to be underconfident), the estimated accuracy of the set as whole will be accurate. This is useful in practice: if the confidence score is biased (e.g., it consistently over-estimates accuracy), then the estimated accuracy of the unlabeled data will systematically differ from its true accuracy. If the score is unbiased, however, then it is useful for evaluating whether the predicted agreement between classifier-produced and manually-produced annotations is sufficient for a given application. To investigate this, we consider the mean signed difference (MSD; see **Methods: Evaluating confidence score calibration**), which quantifies the difference between the predicted accuracy of all predictions in the test set and the actual accuracy of the test set. As shown in **Figure 5C,F**, the temperature scaling-based confidence score has a lower MSD than the softmax-based one, demonstrating that confidence scores derived from temperature scaling are less (positively) biased. While the softmax score consistently overestimates the average accuracy of its predictions by approximately 6-8 percent regardless of training proportion, temperature scaling generally is generally overconfident by only 1-2 percent.

### Uncertainty-based review reduces correction time

Having established the high correspondence between clip confidence score and clip accuracy, we investigate how well our confidence-based review system leverages those confidence scores to reduce the time it takes to review and correct classifier-produced labels. A viable confidence measure would allow clips with a lower confidence score (i.e., lower predicted accuracy) to be preferentially reviewed over those with a higher confidence score, decreasing the manual review time required to obtain acceptably high-quality annotations. Rather than reviewing all the classifier-produced labels, the user could instead review only a portion with the lowest accuracy (see **Methods: Confidence-based review**). We provide an example of this process in practice in **Figure 5G**, which simulates the relationship between the proportion of test video reviewed and the overall accuracy of the labels in the test set. If no video is reviewed, the average accuracy of the test set is the agreement between the classifier produced labels and the ground truth annotations. If one then begins to review and correct videos, the total accuracy increases, since we assume that incorrect classifier-produced labels are corrected. If videos are selected randomly, the relationship between the proportion of the test set reviewed and the test set accuracy is approximately linear – if each video selected is equally likely to have the same number of incorrect labels, then the increase in overall accuracy from correcting those labels is the same for all videos.

If, however, one sorts by confidence measure and reviews the lowest confidence clips first, then, ideally, the subset of videos reviewed will tend to be those with relatively lower accuracy than those not reviewed. The upper bound on the performance of the confidence-based review is a review where the clips are sorted by their actual accuracy (which is what the confidence score approximates). While this is unknown in practice (since the data being reviewed are unlabeled) we simulate it here to provide an upper bound for the performance of the confidence-based review. To compare the performance of the confidence-based review across labeled data proportions, we calculate a metric called “review efficiency” for each split of the data, which expresses the performance of the confidence score bounded by the best (optimal selection, review efficiency of 1) and worst (random selection, review efficiency of 0) possible performance (see **Methods: Evaluating review efficiency**). As shown in **Figure 5H,I,** as the proportion of data labeled increases, both confidence scores become closer to optimal in sorting videos for review. The softmax- and temperature scaling-based scores perform approximately the same.

### Non-technical GUI and flexible codebase improves usability for users with varied technical experience and computational resources

While evaluate our method here using fully annotated datasets, the central purpose of this work is to improve the annotation of behavior in experimental settings. For this reason, we release the entire system as a MATLAB toolbox designed to suit the needs of users with varying technical experience and access to computational resources. The toolbox includes example projects, user-friendly documentation, and GUI interfaces for defining the behavior set of interest (**Fig. 6A**) and conducting manual annotation and confidence-based review (**Fig. 6B**). The annotation GUI is particularly important to packaging this method in a way that is useful for the larger scientific community. For example, we integrate clip-wise annotation by pre-dividing project videos into clips and presenting clips, rather than videos, for users to annotate. In addition, we incorporate the confidence-based review process in a way that is frictionless for users: incomplete (i.e., unreviewed annotations) are shown in a table, with low-confidence clips (and their corresponding confidence scores) appearing at the top so that users can select them for review first. We also include information about the status of the project (e.g., number and duration of videos annotated, video and clip information, etc.) within the GUI. During confidence-based review, we also provide an estimate of 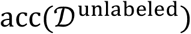 directly, updating it as more annotations are completed. Users can easily load videos, annotate them using the keyboard, add or remove behaviors, and export the results entirely within the GUI.

**Figure 6.**
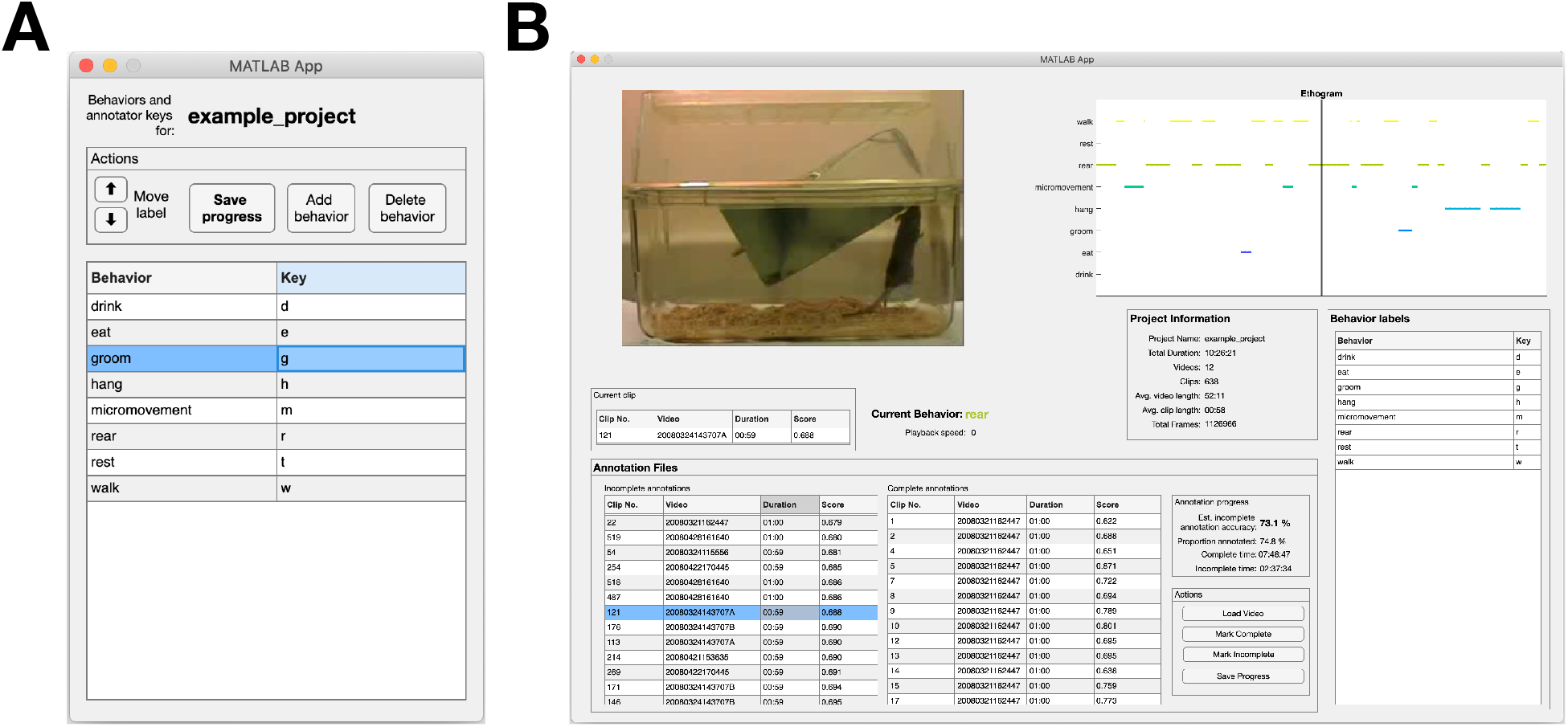
Example usage of the MATLAB apps included in the toolbox. (**A**) GUI for defining the set of behaviors in a dataset. Each behavior label corresponds to a unique keyboard key (“key”), which is used to designate the start and stop of behaviors during manual annotation. (**B**) An example of the annotation GUI used in confidence-based review to correct false classifier-produced predictions. It features tables of the complete (i.e., human annotated or reviewed) and unreviewed (i.e., classifier-annotated) clips in the project. During review, the tables include a confidence score for each clip (“score”) as well as an estimated overall accuracy for all unannotated data. Users select clips to review from the annotation tables, which are then shown in the video viewer box (top left) along with their predicted labels. Users create or correct the labels of the behaviors appearing the video, with both annotation and video playback controlled via keyboard. Behaviors and their corresponding keystrokes are shown in the “Behavior Labels” panel. After completing the annotation of each clip, users press the “Mark Complete” button to save their progress.

## Discussion

In this work we present a method for the automatic annotation of laboratory animal behavior from video. Our classifier produces high accuracy annotations, rivaling or surpassing human-level agreement, while requiring relatively little human annotation time, and performs well across behaviors of varying incidence and timescale. Our confidence scores accurately predict accuracy and are useful in reducing the time required for human annotators to review and correct classifier-produced annotations. Finally, we provide a non-technical, open-source MATLAB toolbox that allows this method to be employed on novel data by researchers with minimal programming experience.

The primary strength of our method is the classifier’s capacity to generate accurate classifications from raw video frames. By classifying behavior using raw frame information, DeepAction removes the need to annotate keypoints and create hand-crafted features that adequately encapsulate a given animals’ behavioral repertoire. This removes both a tedious aspect of manual annotation (i.e., keypoint annotation in addition to behavioral annotation), and alleviates the need for researchers to construct behavior-encapsulating features, which is both time-consuming and often suboptimal. We also note that the performance of DeepAction surpasses that of approaches developed using hand-crafted features on both datasets analyzed (**Table 1**), indicating that the automated feature extraction approach does not compromise performance.

The base level performance of the classifier has the potential to significantly expedite the behavioral research process. Here, a useful benchmark is to compare the accuracy of the classifier (defined as the agreement between classifier-produced labels and the primary set of annotations; see **Methods: Inter-observer reliability**) to the agreement between independent annotators (agreement between the primary set of annotations and a second, independent set used to evaluate inter-observer reliability). In our analysis, we find that the classifier requires only 18 percent of data be annotated to surpass the agreement (71.6 percent) between human annotators on the home-cage dataset (see **Fig. 2A**). Given that the home-cage dataset took 264 hours to manually annotate^7^, if human-level agreement is defined as the threshold for acceptable annotations, our method would reduce this time to 47 hours. Similarly, DeepAction surpasses human-level agreement (69.7 percent) on the CRIM13 dataset with 25 percent of data annotated (see **Fig. 2E**). Since CRIM13 took 350 hours to annotate^6^, using our method instead of manual annotation would have reduced this time to 88 hours, while maintaining annotation quality at the level of human annotators.

Our confidence-scoring system is important for two reasons. The first is a modest increase in review efficiency – if one is to manually review and check some number of automatically-generated behavioral labels, selecting those with the lowest confidence scores is preferable to doing so randomly. We show that this is true across training dataset sizes, and that review using confidence scores becomes closer to optimal as more data is annotated. The second, and perhaps more important, reason is that the temperature scaling-based confidence score generates an accurate estimate of the overall agreement between classifier- and human-produced labels on unlabeled data (i.e., where the “human-produced labels” are unknown). This means that researchers could annotate data until the estimated accuracy of the unlabeled data reached a given threshold of acceptable agreement for their given behavioral analysis, and then export the automated annotations without having to review and correct them.

Our tool has several practical advantages. First is accessibility to those without technical background; in addition to releasing a GUI for annotation and confidence-based review, we provide a repository with documentation and details example projects. Second is adaptability; in our GitHub release we provide additional pretrained CNNs (e.g., ResNet50 and Inception ResNetv2) with which potentially more useful features could be extracted, a computationally faster optical flow algorithm^21^, and options to parallelize a number of the computationally-intensive project functions (e.g., temporal frame generation and feature extraction). A final advantage is modularity: users can use the classification portion of the workflow without the review component, the annotator can for its interface alone, etc.

Though the toolbox presented here represents a significant advancement as compared to entirely manual annotation, there are several avenues for further exploration and potential improvement. While our clip selection process demonstrates superior performance to whole-video annotation, in the results here we select clips randomly. In practice, the confidence-based review system can be used to iteratively train the classifier (**Fig. 1A**), where low-confidence clips are reviewed, corrected, and used to re-train the classifier (though we do not explore whether this is preferable to random selection here). An alternative approach would be to adapt methods used in video summarization to cluster video clips by their similarity, and then select the subset of clips best representative of the overall dataset^22,23^. While our classifier is based on a LSTM with bidirectional layers, it is possible that alternate architectures would demonstrate superior performance^11,24^. Relatedly, the classifier described here assumes behaviors are mutually exclusive; that is, none of the behaviors can occur at the same time. However, for datasets in which this is not the case, the cross-entropy loss function used here could easily be adjusted to allow for co-occurring behaviors. A final avenue for further exploration is our approach to calculating confidence scores. Though our system is already close to optimal, given enough training data (**Fig. 5H,I**), there are a number of density-based metrics^25^ or those that utilize Bayesian dropout^26^ that might provide superior performance to the temperature scaling-based one employ here.

## Methods

### Datasets

Given that rodents are widely used in behavioral research, and mice are the most studied rodents^27^, we chose two publicly-available datasets featuring mice engaging in a range of behaviors. The first dataset, referred to as the “home-cage dataset,” was collected by Jhuang, et al. ^7^ and features 12 videos (approximately 10.5 hours and 1.13 million frames in total) of singly housed mice in their home cages, recorded from the side view. Video resolution is 320 × 240 pixels. The authors annotate each video in full and identify eight mutually exclusive behaviors (**Fig. S1A**) of varying incidence (**Fig. 3A**). This dataset allows us to benchmark our approach against existing methods, allows us to evaluate our method on a common use-case, and is relatively well-balanced in terms of the incidence of each behavior.

The second dataset used is the Caltech Resident-Intruder Mouse dataset (CRIM13), collected by Burgos-Artizzu, et al. ^6^. In consists of 237 pairs of videos, recorded from synchronized top- and side-view cameras, at 25 frames per second and an 8-bit pixel depth. Videos are approximately 10 minutes long, and the authors label 13 mutually exclusive actions (**Fig. S1B**). Of these actions, 12 are social behaviors, and the remaining action is the category “other,” which denotes periods where no behavior of interest occurs^6^. This dataset features a number of challenges absent from the Jhuang, et al. ^7^ dataset. In addition to including social behavior (in contrast to the home-cage dataset, which features singly-house mice), it presents two algorithmic challenges. First, videos are recorded using a pair of synchronized cameras. This allows us to test multiple-camera integration functionality (see **Methods: Feature extraction**), to evaluate classifier performance using features from multiple cameras. And second, it is highly unbalanced, with a slight majority of all annotations being the category “other” (periods during which no social behavior occurred; **Fig. 3D**).

### Inter-observer reliability

Both datasets include a set of annotations performed by two groups of annotators. The primary set of annotations was produced by the first group of annotators an includes all video in the dataset. The secondary set of annotations was performed by a second, independent set of annotators on a subset of videos. We use the primary set of annotations to train and evaluate our method, and the secondary set to establish inter-observer reliability; that is, how much two, independent human annotator’s annotations can be expected to differ. Given this, classifier-produced labels can be most precisely interpreted as the predicted behavior *if the video was annotated by the first group of annotators.* This distinction becomes important because we benchmark the accuracy of our method (i.e., the agreement between the classifier’s predictions and the primary set of annotations) relative to the inter-observer agreement (i.e., the agreement between the first and second group of annotators, on the subset of video labeled by both groups). So, for example, when we note that our model achieves accuracy “above human agreement,” we mean that our classifier predicts the labels from the first human annotator group better than the second human annotator group does. In the case of the home-cage dataset, the agreement between the primary and secondary sets was 78.3 percent, compared on a 1.6h subset of all dataset video^7^. For CRIM13, agreement was 69.7 percent, evaluated on a random selection of 12 videos^6^.

### Simulating labeled data

To simulate our approach’s performance with varying amount of training data, in our primary analyses we train the classifier using the following amounts of labeling:

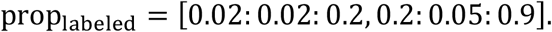

That is, we use a proportion of all data, prop_labeled_, to construct our training and validation sets (i.e., 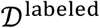), and the remaining 1 – prop_labeled_ data to create our test set, 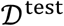 (**Fig. 1B,D**). We use an increment of 0.02 for low training proportions (up to 0.20), because that is when we see the greatest change relative to a small change in added training data (**Fig. 2A,E**). We increment values from 0.25 to 0.90 by 0.05. This gives us a set of 24 training proportions per analysis. Additionally, for each training proportion, unless otherwise noted, we evaluate the model on 10 random splits of the data. In our main analyses, we use a clip length of one minute for both datasets.

### Comparison with existing methods

When comparing our model to existing methods, we employ *k*-fold validation instead of evaluating on random splits of the data. In the case of the home-cage dataset, the existent methods cited employ a “leave one out” approach – using 11 of the 12 videos to train their methods, and the remaining video to test it. In our approach, however, we rely on splitting the data into clips, so instead we use 12-fold cross-validation in which where we randomly split the dataset clips into 12 folds and then employ cross-validation on the clips, rather than entire videos. In evaluating their approach’s performance on the CRIM13 dataset, Burgos-Artizzu, et al. ^6^ selected 104 videos for training and 133 for testing, meaning that they trained their program on 44 percent of the data, and tested it on 56 percent. Here, we evaluate our method relative to theirs using two-fold cross validation (50 percent test and 50 percent train split) to retain similar levels of training data.

### Frame extraction

To generate spatial frames, we extract raw video frames from each video file. Rather than save each image as an image file in a directory, we save the entire sequence of images corresponding to a single video to a sequence file, using the implementation provided by Dollár ^28^ with JPG compression. This has the advantage of making the video frames easier to transfer between file systems and readable on any operating system (which is useful for users running the toolbox on high performance computing clusters). To generate the temporal component, we use the TV-L1 algorithm^19,29^, which shows superior performance to alternate optical flow algorithms^15^, to calculate the dense optical flow between pairs of sequential video frames and represent it visually via the MATLAB implementation by Cun ^30^. In the visual representation of optical flow fields, hue and brightness of a pixel represent the orientation and magnitude of that pixel’s motion between sequential frames. By representing motion information of the video as a set of images, we can use a similar feature extraction method for both spatial and temporal frames. Just as the features derived from the spatial images represent the spatial information in the video, the features derived from the temporal images should provide a representation of the motion information in the video.

### Feature extraction

We utilize the pretrained ResNet18 convolutional neural network (CNN) to extract high-level features from the spatial and temporal video frames. Often used in image processing applications, CNNs consist of a series of layers that take an image as an input and generate an output based on the content of that image. Intuitively, classification CNNs can be broken down into two components: feature extraction and classification. In the feature extraction component, the network uses a series of layers to extract increasingly complex features from the image. In the classification component, the network uses the highest-level features to generate a final classification for the image (e.g., “dog” or “cat”). In the case of pretrained CNNs, the network learns to extract important features from the input image through training – by generating predictions for a set of images for which the ground truth is known, and then modifying the network based on the deviation of the predicted classification from the true classification, the network learns which features in the image are important in discriminating one object class from another. In pretrained CNNs, such as the ResNet18, which was trained to categorize millions of images from the ImageNet database into one thousand distinct classes^31^, early layers detect generic features (e.g., edges, textures, and simple patterns) and later layers represent image data more abstractly^32^.

Here, we leverage transfer learning – where a network trained for one context is used in another – to extract a low-dimensional representation of the data in the spatial and temporal video frames. The idea is that, since the ResNet18 is trained on a large, general object dataset, the generality of the network allows us to obtain an abstract representation of the salient visual features in the underlying video by extracting activations from the later layers of the network in response to a completely different set of images (in this case, laboratory video of animal behavior). To extract features from the ResNet18 network for a given image, we input the image into the network and record the response (“activations”) from a specified layer of the network. In this work, we chose to extract activations from the global average pooling layer (“pool5” in MATLAB) of the ResNet18, close to the end of the network (to obtain high-level feature representations). This generates a feature vector of length 512, representing high level CNN features for each image.

By default, the ResNet18 accepts input images of size [224,224, 3] (i.e., images with a width and height of 224 pixels and three color channels), so we preprocess frames by first resizing them to a width and height of 224 pixels. In the case of spatial frames, the resized images are input directly into the unmodified network. For temporal frames, however, rather than inputting frames into the network individually, we “stack” each input frame to the CNN with the five frames preceding it and the five frames following it, resulting in an input size of [224, 224, 33]. This approach allows the network to extract features with longer-term motion information and has been shown to improve discriminative performance^14,18^. We select a stack size of 11 based on the findings from Simonyan and Zisserman ^18^. By default, the ResNet18 network only accepts inputs of size [224,224,3], so to modify it so that it accepts inputs of size [224,224,33] we replicate the weights of the first convolutional layer (normally three channels) 11 times. This allows the modified “flow ResNet18” to accept stacks of images as inputs, while retaining the pretrained weights to extract salient image features.

After spatial features and temporal features have been separately extracted from the spatial and temporal frames, respectively, we combine them to produce the *spatiotemporal* features that will be used to train the classifier (**Fig. 1E**). To do so, we simply concatenate the spatial and temporal features for each frame. That is, for a given segment of video with *n* frames, the initial spatiotemporal features are a matrix of size [*n,* 512 × 2] = [*n,* 1024], where 512 represents the dimensionality of the features extracted from the ResNet18. If multiple synchronized cameras are used (as is the case in one of our benchmark datasets), we employ the same process, concatenating the spatial and temporal features for each frame *and each camera.* In the case of two cameras, for example, this implies the initial spatiotemporal features is a matrix of size [n, 512 × 2 × 2] = [n, 2048]. To decrease training time, memory requirements, and improve performance^33,34^, we utilize dimensionality reduction to decrease the size of the *initial* spatiotemporal features to generate *final* spatiotemporal features of size [n, 512]. We selected reconstruction independent component analysis^35,36^ as our dimensionality reduction method, which creates a linear transformation by minimizing an objective function that balances the independence of output features with the capacity to reconstruct input features from output features.

### Classifier architecture

The labeled and unlabeled data consist of a set of clips, generated from project video, which the classifier uses to predict behavior. Clips in 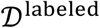 and 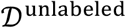 are both constituted of a segment of video and a corresponding array of spatiotemporal features extracted from that video. Clips in 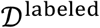 also include an accompanying set of manual annotations (**Fig. 1B**). For a given clip in 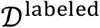 with *n*_labeled_ frames, the classifier takes a [*n*_labeled_, 512]-dimensional vector of spatiotemporal features (**Fig. 1E**) and a one-dimensional array of *n*_labeled_ manually-produced labels (e.g., “eat,”“drink,” etc.) as inputs, and learns to predict the *n*_labeled_ labels from the features. After training, for a given clip in 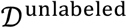 with *n*_unlabeled_ frames, the classifier takes as an input a [*n*_unlabeled_, 512]-dimensional vector of spatiotemporal features and outputs a set of *n*_unlabeled_ behavioral labels, corresponding to the predicted behavior in each of the *n*_unlabeled_ frames. To implement this transformation from features to labels, we rely on recurrent neural networks (RNNs). Prior to inputting clips into the RNN, we further divide them into shorter “sequences,” corresponding to 15 seconds of video to reduce overfitting^37^ and sequence padding^38^. Unlike traditional neural networks, recurrent neural networks contain cyclical connections which allows information to persist over time, enabling them to learn dependencies in sequential data^39^. Given that predicting behavior accurately requires the integration of information over time (i.e., annotators generally must view more than one frame to classify most behavior, since behaviors are often distinguished by movement over time), this persistence is critical.

We opt for a long short-term memory (LSTM) network with bidirectional LSTM layers (BiLSTM) as the core of our classification model. LSTMs are better able to learn long-term dependencies in data than traditional RNNs in practice^40,41^, and the use of bidirectional layers allows the network to process information in both temporal directions^42^ (i.e., forward and backward in time, rather than forward only in the case of a traditional LSTM layer). As shown in **Figure S4**, our network’s architecture begins with a sequence input layer, which accepts a two-dimensional array corresponding to spatiotemporal video features (with one row per frame and one column per feature). We then apply two BiLSTM layers, which increases model complexity and allows the model to learn more abstract relationships between input sequences and correct output labels^43^. To reduce the likelihood of model overfitting, we use a dropout layer after each BiLSTM layer, which randomly sets some proportion of input units (here, 50 percent) to 0, which reduces overfitting by curbing the power of any individual neuron to generate the output^44^. The second dropout layer is followed by a fully-connected layer with an output size of [*n*, *K*], where *K* is the number of classes and *n* is the number of frames in the input clip. The softmax layer then normalizes the fully-connected layer’s output into a set of class probabilities with shape [*n*, *K*], where the sum of each row is 1 and the softmax probability of class *k* in frame *j* is given by the entry *jk*. Following the softmax layer, the sequence-to-sequence classification layer generates a one-dimensional categorical array of *n* labels corresponding to the behavior with the highest softmax probability in each frame. We select cross-entropy loss for *K* mutually exclusive classes^45^ as our loss function, since the behaviors in both datasets are mutually exclusive. All classifiers were trained using a single Nvidia Tesla K80 GPU running on the Dartmouth College high performance computing cluster.

### Classifier training

In this analysis, use the hyperparameters specified in **Table 2** when training the network. To avoid overfitting, we select 20 percent of 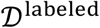 to use in our validation set (i.e., prop_*train*_ = 0.20; see **Fig. 1C**). We then evaluate the network on this validation set every epoch (where “epoch” is defined as a single pass of the entire training set through the network) and record its cross-entropy loss. If the loss on the validation set after a given epoch is larger than or equal to the smallest previous loss on the validation set more than twice, training terminates.

**Table 2.** Default hyperparameters. The maximum number of epochs the network can be trained for is 16. The cross-entropy loss of the validation dataset is calculated for each epoch, and if this value is less than the prior minimum validation loss for more than two epochs, training terminates. The initial learning rate is 0.001, and every four epochs the learning rate drops by a factor of ten. We use a minibatch size of eight to minimize deficits in generalizability that could occur at larger values^46^.

| Hyperparameter | Value |
| --- | --- |
| Maximum epochs | 16 |
| Validation frequency (per epoch) | 1 |
| Validation patience | 2 |
| Initial learning rate | 0.001 |
| Learning rate drop period | 4 |
| Learning rate drop factor | 0.1 |
| Minibatch size | 8 |

### Classifier evaluation

To evaluate the classifier, we consider its performance on the test set, 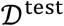 (**Fig. 1B,D**). For each clip, the classifier outputs a set of predicted labels for each frame, corresponding to the predicted behavior in that frame. In evaluating the classifier, we are interested in how closely these predicted labels match the true ones. We first consider overall prediction accuracy. We let correct denote the number labels in which the network’s prediction is the same as the true label and incorrect the number of labels in which the network’s prediction is not the same as the true label. Then accuracy can be quantified as the following proportion:

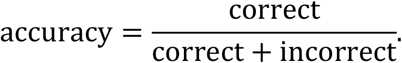

Next, we consider the performance of the network by behavior. To do so, we let TP_*k*_ denote the number of true positives (predicted class *k* and true class *k*), FP_*k*_ the number of false positives (predicted class *k*, but true label not class *k*), and FN_*k*_ the number of false negatives (true class *k*, predicted not class *k*) for class *k* (where *k* is between 1 and the total number of classes, *K*).

We then calculate the precision, recall, and F1 score for each label^11,47^, where the precision and recall for class *k* are defined as follows:

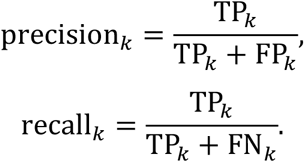

Precision is the proportion of correct predictions out of all cases in which the *predicted class* is class *k*. Recall, meanwhile, denotes the proportion of correct predictions out of all the cases in which the *true class* is class *k*. From the precision and recall, we calculate the F1 score for class *k*. The F1 score is the harmonic mean of precision and recall, where a high F1 score indicates both high precision and recall, and deficits in either decrease it:

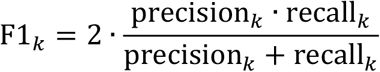

After calculating the F1 score for each class, we calculate the average F1 score, F1_all_ as follows:

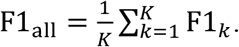

### Confidence score definition

For each input clip, the classifier returns a set of predicted annotations corresponding to the predicted behavior (e.g., “walk,” “drink,” “rest,” etc.) occurring in each frame of that clip. We denote the set of classifier-predicted labels for clip number *i*, clip_*i*_, as 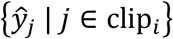. Each clip also has a set of “true” labels, corresponding to those that would be produced if the clip was manually annotated. In the case of the labeled data, the true labels are known (and used to train the classifier). In the case of unlabeled data, they are not known (prior to manual review). We denote the set of true labels for clip_*i*_ as {*y_j_* | *j* ∈ clip_*i*_}. For each frame in a clip, in addition to outputting a prediction for the behavior occurring in that frame, we also generate an estimate of how likely that frame’s classifier-assigned label is correct. That is, for each clip, we generate a set of predicted probabilities 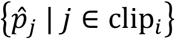 such that 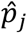 denotes the estimated likelihood that 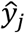 is equal to *y_j_*. In an optimal classifier, 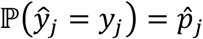. That is, 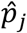 is an estimate of the probability the classification is correct; and, in an optimal confidence-scorer, the estimated probability the classification is correct will be the ground truth likelihood the classification is correct^48^.

Now that we have established an estimated probability that a given *frame* in a clip is correct, we extend the confidence score to an entire clip. As in training data annotation, the review process is conducted at the level of an entire clip, not individual video frames. That is, even if there are a handful of frames in a clip that the classifier is relatively unconfident about, we assume that a human reviewer would need to see the entire clip to have enough context to accurately correct any misclassified frames. Since 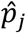 is the estimated probability a given frame *j* is correct, it follows that the average 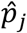 for *j* ∈ clip_*i*_ is the estimated probability a randomly selected frame in clip_*i*_ is correct. We define this quantity to be the clip confidence score; formally, 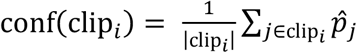, where conf(clip_*i*_) is the clip confidence score of clip_*i*_ and |clip_*i*_| is the number of frames in clip_*i*_. We then consider that accuracy is the true probability a randomly selected frame in clip_*i*_ is correct by definition. That is, 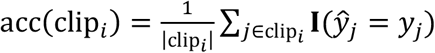, where acc(clip_*i*_) is the accuracy of clip_*i*_ and **I** is the indicator function. In the case of an optimal confidence score, we’ll have that conf(clip_*i*_) = acc(clip_*i*_). If we compare conf(clip_*i*_) with acc(clip_*i*_) on our test data, we can establish how well the confidence score can be expected to perform when the ground truth accuracy, acc(clip_*i*_), is unknown. In **Methods: Confidence score calculation**, we discuss our approach for obtaining 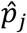, after which finding clip-wise confidence scores is trivial.

### Confidence score calculation

Here, we first examine how to calculate the frame-wise confidence score 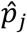. To do so, we consider the classifier structure (**Fig. S4**) in more detail. In particular, we focus on the last three layers: the fully-connected layer, the softmax layer, and the classification layer. To generate a classification for a given frame, the softmax layer takes in a logits vector from the fully-connected layer. This logits vector represents the raw (unnormalized) predictions of the model. The softmax layer then normalizes these predictions into a set of probabilities, where each probability is proportional to the exponential of the input. That is, given *K* classes, the *K*-dimensional vector from the fully-connected layer is normalized to a set of probabilities, representing the probability of each class. The class with the highest probability is then returned as the network’s predicted label (e.g., “eat” or “walk”) for that frame. We can then interpret this probability as a confidence score derived from the softmax function^48^. Formally, if we let logits vector ***z**_j_* represent the output from the fully-connected layer corresponding to frame *j*, the softmax-estimated probability the predicted label of frame *j* is correct is 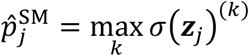, where *σ* is the softmax function. We refer to this confidence score as the “max softmax score,” since it is derived from the maximum softmax probability.

One of the challenges with using the max softmax probability as a confidence score, however, is that it is often poorly scaled. Ideally, estimated accuracy for a prediction would closely match its actual expected accuracy, but in practice the softmax function tends to be “overconfident”^48^. That is, 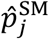 tends to be larger than 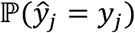. To generate a more well-calibrated confidence score (i.e., one in which 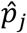 is closer to 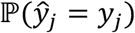, we use an approach called temperature scaling. Temperature scaling uses a learned parameter *T* (where *T* > 1 indicates decreased confidence and *T* < 1 increased confidence) to rescale class probabilities so that the confidence score more closely matches the true accuracy of a prediction^49^. We define the temperature scaling-based confidence for frame *j* as 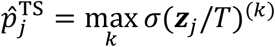, where *T* is selected to minimize the negative log likelihood on the validation set. Now that we have established the process for generating a frame-wise confidence score, we can generate the clip-wise confidence score that is used in the confidence-based review. As previously described, for clip_*i*_ this is simply 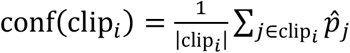, where 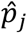 is either generated via the softmax function 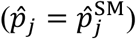 or temperature scaling 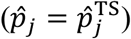.

### Confidence-based review

Now that we have generated a confidence score for a given clip, we use it in two ways. First, recall that one of the purposes of the confidence-based review is to estimate the accuracy of the unlabeled data, 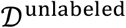. If, for example, a user decided that an accuracy of 80 percent was acceptable for their given behavior analysis application (i.e., 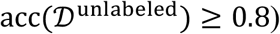), then given an acceptably reliable confidence score, unlabeled data for which 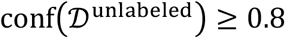 would be sufficient for export and use in their given analysis without manual review. Before obtaining an estimate for 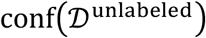, we first consider that the true (unknown) accuracy of the annotations in 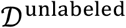 is the weighted sum of the accuracies of the clips in 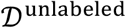, where weight is determined by the number of frames in each clip. Formally, we can express the accuracy of 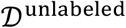 as:

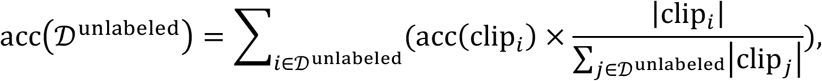

where 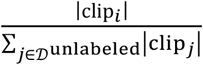 weights the accuracy of acc(clip_*i*_) by the number of frames in clip_*i*_ (i.e., |clip_*i*_|) relative to the total number of clips (i.e., 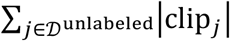). We then estimate the accuracy of the unlabeled data by substituting the known conf(clip_*i*_) for the unknown acc(clip_*i*_):

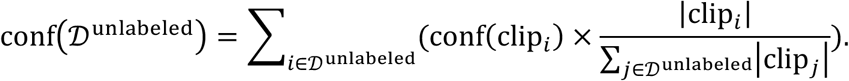

In this way, 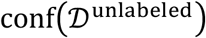 represents the approximate accuracy of the classifier on unlabeled data. If the confidence score functions well, then 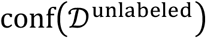 will closely match 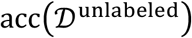.

Next, we consider the confidence-based review. In this component of the workflow, user can review and correct labels automatically generated by the classifier for 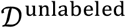. A naïve approach would be to review all the video clips contained in 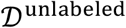. While this would indeed ensure all the labels produced by the classifier are correct, if 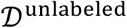 is large it can prove quite time-consuming. So instead, we leverage confidence scores to allow users to only annotate the subset of clips with relatively low confidence scores (i.e., relatively low predicted accuracy), for which review is most productive, while omitting those with relatively high confidence scores.

If a user reviews only a portion of the clips, it should be the portion with the lowest accuracy, for which correction is the most important. To express this formally, consider an ordered sequence of the *n* clips in 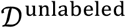, (clip_1_, clip_2_,…, clip_n_), sorted in ascending order by accuracy (i.e., acc(clip_*i*_) ≤ acc(clip_*j*_), for *i* < *j* and all *i*,*j* ≤ *n*). If we review only *k* of the *n* clips, where *k* ≤ *n*, we are best off reviewing clips clip_1_,clip_2_,…,clip_*k*_ from the list since they have the lowest accuracy. For unlabeled data, however, recall that we can’t precisely sort clips by accuracy, since without ground truth annotations acc(clip_*i*_) is unknown. However, since conf(clip_*i*_) approximates acc(clip_*i*_), we can instead sort unlabeled clips by their (known) confidence scores, and then select the clips with the lowest confidence scores to review first. This forms the basis of the confidence-based review. Given a set of clips in 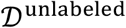, we simply create a sequence of clips (clip_1_, clip_2_,…, clip_*n*_) sorted by confidence score (i.e., such that conf(clip_*i*_) ≤ conf(clip_*j*_), for all *i* < *j*) and then have users review clips in ascending order. If the confidence score is an effective estimate of the clip accuracies, sorting based on confidence score will approximate sorting by accuracy.

### Evaluating confidence score calibration

To examine the relationship between confidence scores and accuracy, we first consider the relationship between individual clips’ predicted accuracy (as derived from confidence cores) and actual accuracy. The prediction error (PE) for a given clip is defined as the signed difference between its predicted accuracy and its actual accuracy. For clip_*i*_, the PE is then PE(clip_*i*_) = conf(clip_*i*_) – acc(clip_*i*_). Positive values indicate an overconfident score, and negative value and underconfident one. The absolute error (AE) is the magnitude of the prediction error and is defined as AE(clip_*i*_) = |PE(clip_*i*_)|. The AE is always positive, with a higher AE(clip_*i*_) indicating a greater absolute deviation between conf(clip_*i*_) and acc(clip_*i*_).

While PE and AE are defined for a single clip, we also consider the mean absolute error and mean prediction error across all the clips in 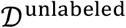. Here, we let clip_1_, clip_2_,…, clip_*n*_ denote a set of *n* clips. The mean absolute error (MAE) is defined as 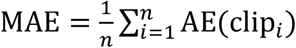. MAE expresses the average magnitude of the difference between predicted accuracy and actual accuracy for a randomly selected clip in the set. So, for example, if MAE = 0.1, then a randomly selected clip’s confidence score will differ from its accuracy score by about 10 percent, in expectation. The mean signed difference (MSD), meanwhile, is defined as 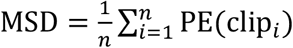. MSD expresses the signed difference between the total expected accuracy across clips and the total actual accuracy. So, for example, is MSD = −0.05, then the total estimated accuracy of annotations for the set clip_1_, clip_2_,…, clip_n_ is five percent lower than the true accuracy.

### Evaluating review efficiency

To develop a metric for the performance of the confidence-based review, we first consider a case where a user has generated predicted labels for *n* clips, which have not been manually labeled, and selects *k* of them to review, where *k* ≤ *n*. The remaining *n* – *k* clips are not reviewed and are exported with unrevised classifier-generated labels. Then, for each of the *k* clips the user has selected, he or she reviews the clip and corrects any incorrect classifier-generated labels. In this formation, after reviewing a given clip, that clip’s accuracy (defined as the agreement between a clip’s labels and the labels produced by manual annotation), is 1, since any incorrect classifier-produced labels would have been corrected.

Next, we assume that we have been provided with a *sequence* of *n* clips, 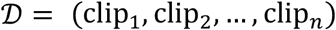, from which we select the first *k* clips in the sequence to review. If we denote 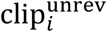 as clip *i* prior to being reviewed, and 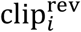 as clip *i* after being reviewed, then we can express the sequence of the first *k* clips after they have been reviewed as 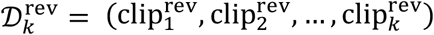. We then express the remaining *n* – *k* clips as the sequence 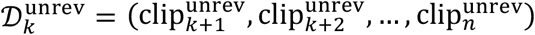. We then consider that the overall accuracy of the sequence of clips, 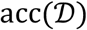, is simply weighted average of the accuracy of the reviewed videos, 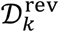, and the unreviewed ones, 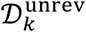, where the weight is a function of the number of frames in each clip.

Formally,

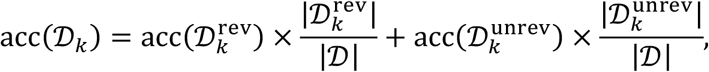

where 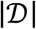 is the total number of video frames in the clips in set 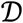 (i.e., 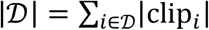). We then consider that, after reviewing and correcting the first *k* clips, the accuracy of each reviewed clip is now 1. That is, 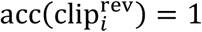 for all 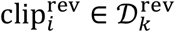. Therefore, the total accuracy of sequence 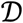, after reviewing the first *k* clips, is

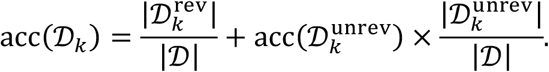

This method for calculating the accuracy of dataset 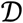 after reviewing the first *k* clips becomes useful for analyzing the performance of the confidence-based review. To see why, we first consider the lower bound on 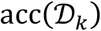. In the worst case, our confidence score will convey no information about the relative accuracies of the clips in 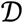. Without a relationship between acc(clip_*i*_) and conf(clip_*i*_), sorting based on confidence score is effectively the same as randomly selecting clips. In this way, we can compare the accuracy after labeling the first *k* clips via confidence-score with the accuracy that *would have been obtained if the first* k *clips were reviewed.* We denote this improvement in accuracy using confidence metric conf as the “improvement over random” and formalize it as 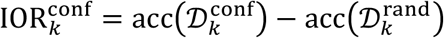, where 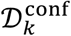 and 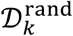 denote dataset 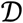 sorted by confidence score and randomly, respectively.

Next, we place an upper bound on IOR_*k*_ by considering the maximum accuracy that 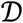 could have after reviewing *k* clips. In the best case, the first *k* clips reviewed would be the *k* clips with the lowest accuracy. Here, since we’re evaluating on 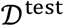, where accuracy is known, we can calculate this. If we let 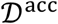 denote the sequence of clips sorted in ascending order by their true accuracy, then the maximum accuracy of 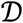 after reviewing *k* clips is 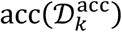. Then, similar to the analysis above, we calculate the improvement of optimal review (i.e., review based on true accuracy) over random review as 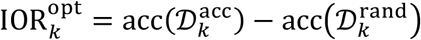. Semantically, 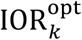 expresses how much higher the accuracy of the test set it after reviewing *k* clips in the optimal order than it would be if clips had been reviewed randomly.

We can then derive a series of global measures for the confidence-based review. While IOR_*k*_ is defined for a single number of clips reviewed, *k*, we look to generate a measure that expresses IOR_*k*_ across a range of *k* values. To do so, we calculate the average improvement over random across the number of clips reviewed, from 0 to the total number, *n*, as follows:

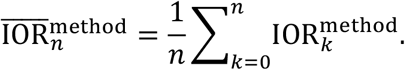

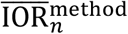 the mean improvement over random of method method over *n* clips. After calculating 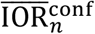 and 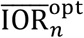 (i.e., 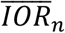 for confidence-based and optimal sorting), we can generate a final measure for the review efficiency by expressing the average improvement of confidence score conf over random relative to the maximum possible improvement over random (optimal review):

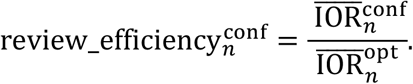

This metric expresses how close review using metric conf is to optimal. If sort order based on conf exactly matches that of sorting by accuracy, 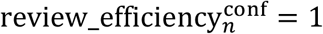. If the sort order is no better than random, 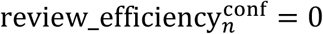.

### Implementation details and code availability

We implement the toolbox in MATLAB version 2020b. The GUI for annotation and confidence-based review is included in the toolbox as a MATLAB application. Figures are produced using Prism9 and OmniGraffle. The entire toolbox, along with example scripts, documentation, and additional implementation details is hosted via a public GitHub repository at: https://github.com/carlwharris/DeepAction. We provide the intermediary data generated for the home-cage dataset (e.g., spatial and temporal frames and features, annotations, etc.) as an example project linked in the GitHub repository. Data produced to generate results for the CRIM13 project is available upon request, but not provided as an example project due to its large file sizes. Full data (i.e., results for each test split in both projects) needed to replicate the results is also available on request.

## Acknowledgments

Supported by National Science Foundation Award #1632738 (PUT), the Neukom Institute for Computational Science at Dartmouth College (Neukom Scholars award to CH and Neukom Post-doctoral Fellowship to KF), and the David C. Hodgson Endowment for Undergraduate Research Award (to CH). Any opinions, findings, and conclusions or recommendations expressed in this material are those of the author(s) and do not necessarily reflect the views of the National Science Foundation.

## Author contributions

CH conceived of the project, developed the approach and software, and analyzed the results (with input from KF). CH wrote the manuscript with input from PT and KF.

## Additional information

Supplementary information is available for this paper. Correspondence and requests for materials should be addressed to the corresponding author.

## Extended data figures

**Figure S1.**
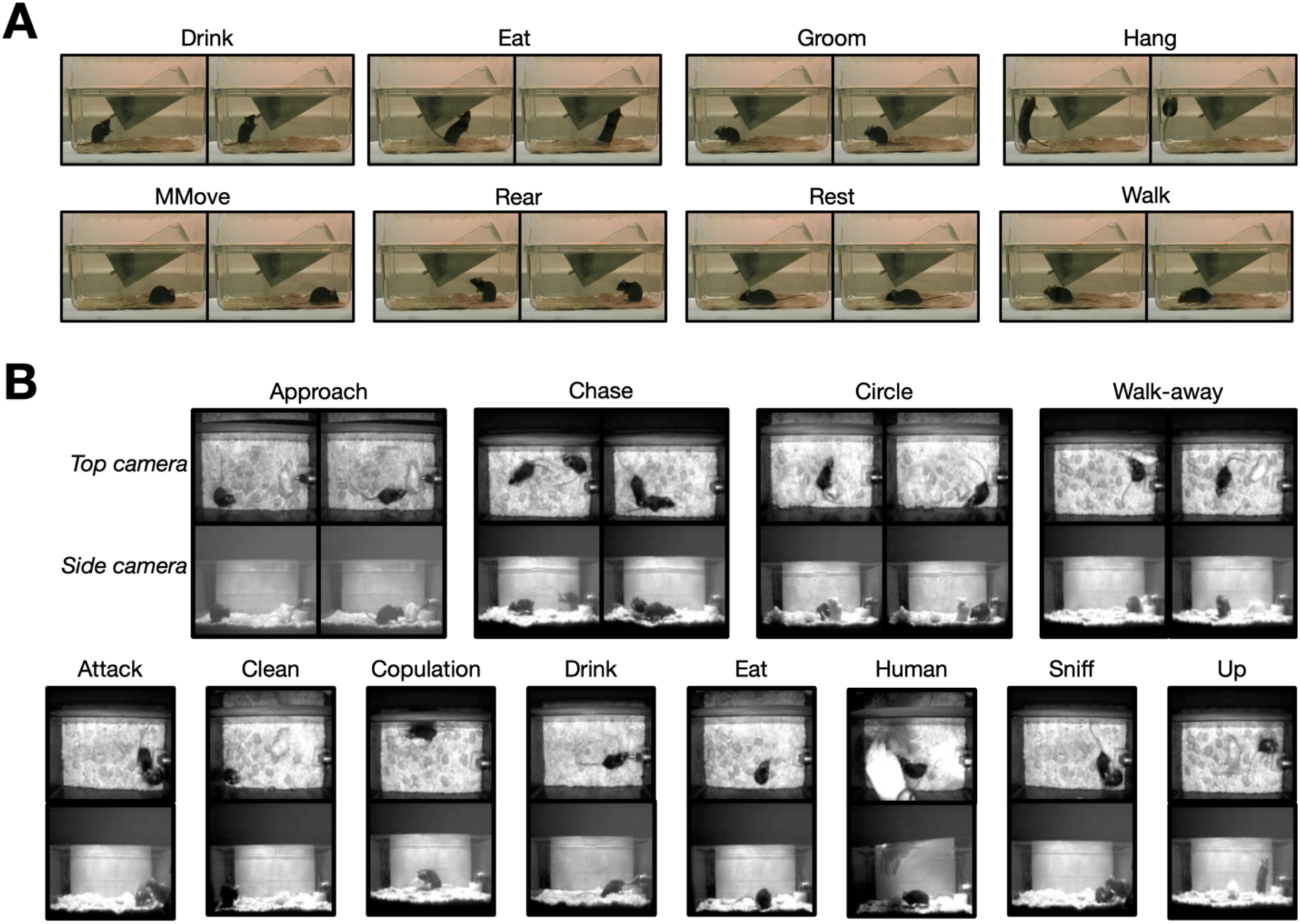
Example behaviors. (**A**) Examples of the eight behaviors included in the home-cage dataset. (**B**) Examples of the 12 social behaviors of interest in the CRIM13 database (“other” not shown). Screenshots in (**B**) from Burgos-Artizzu, et al. ^6^.

**Figure S2.**
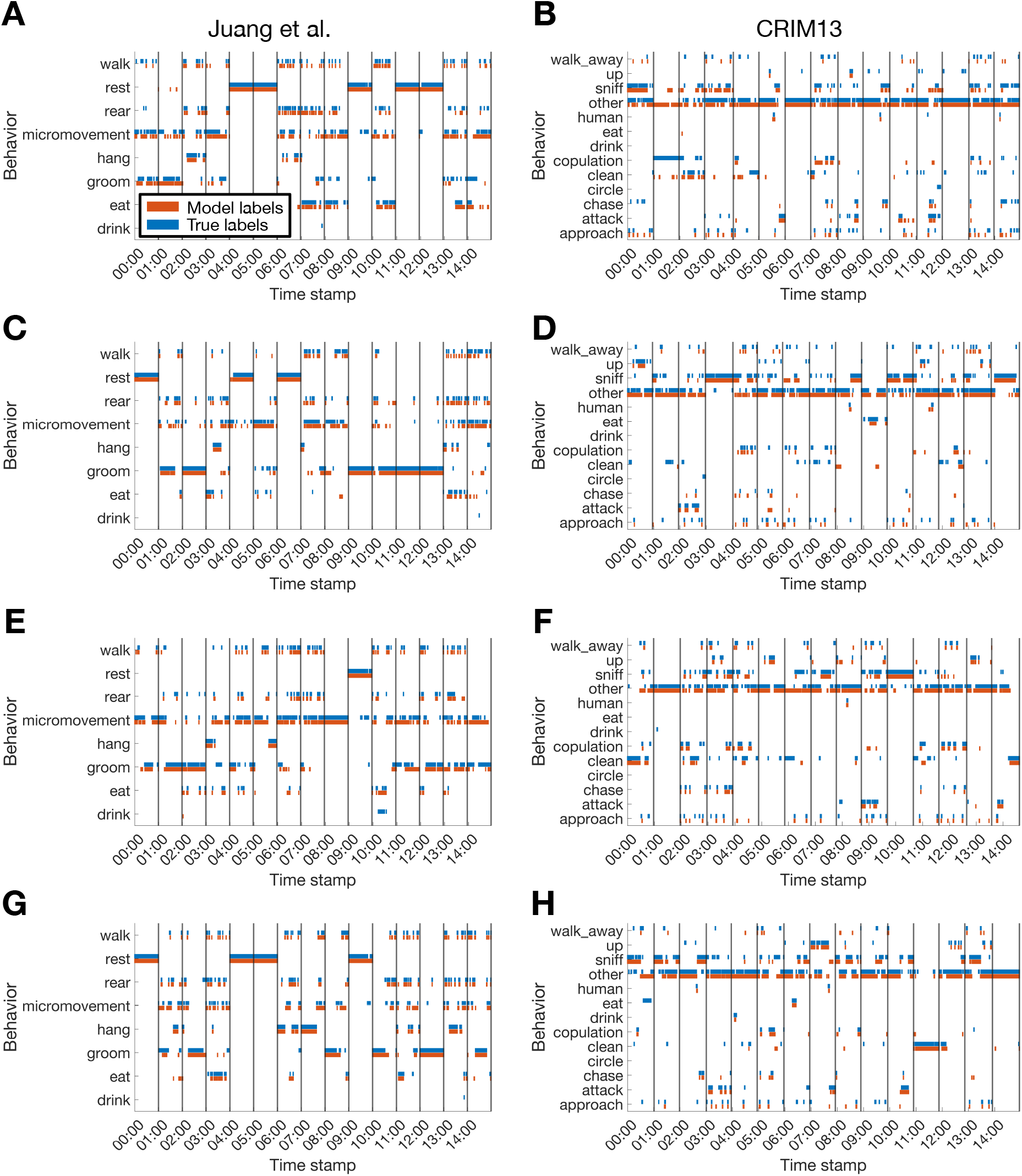
Sample ethograms with different training data proportions. Plotted are ethograms corresponding to 20 minutes of randomly-selected clips with (**A,B**) 10, (**C,D**) 20, (**E,F**) 50, and (**G,H**) 80 percent of data used to train the classifier. Black lines denote the start of a new clip, and the data between the black lines denotes the ethogram corresponding to that clip.

**Figure S3.**
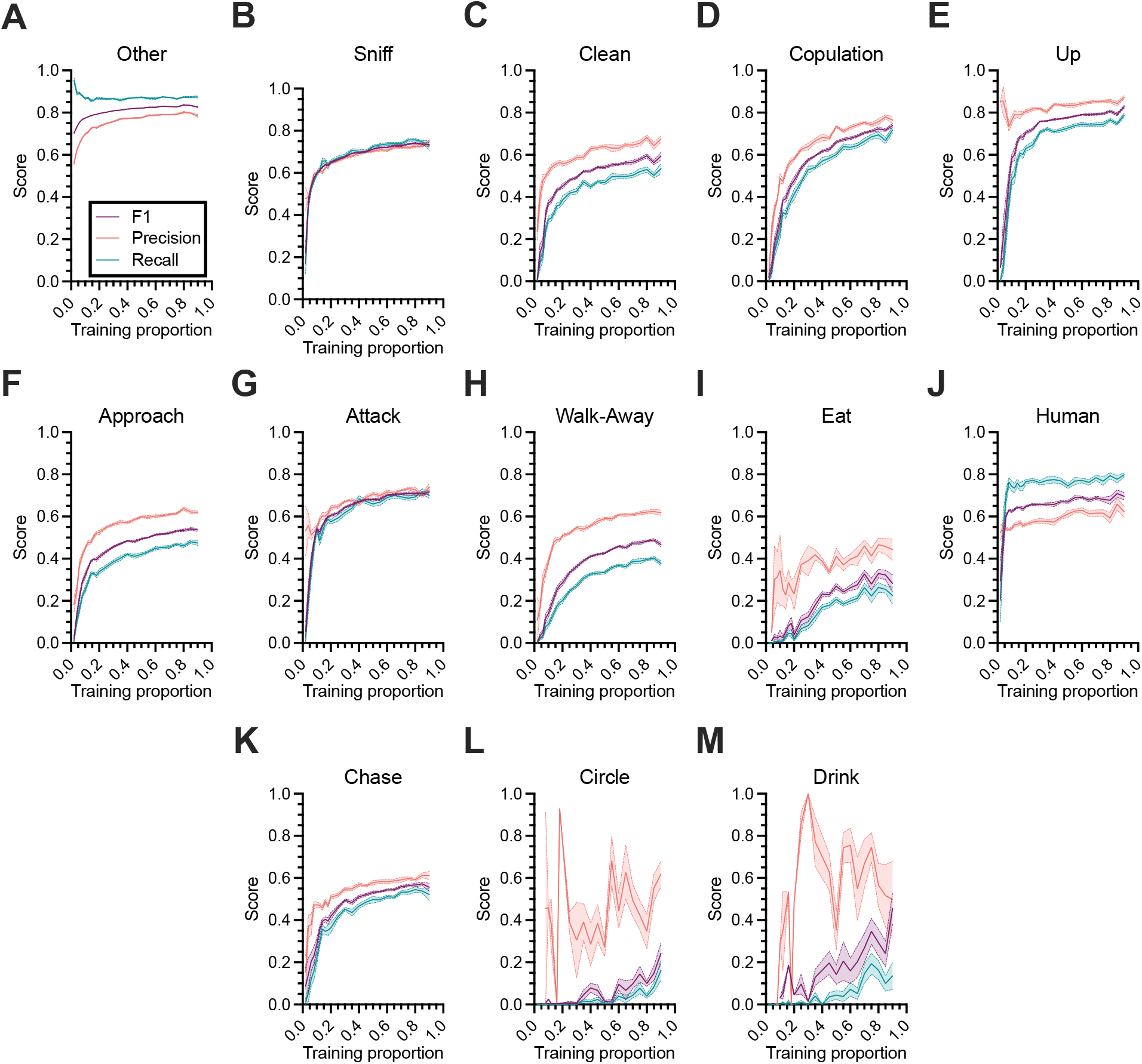
CRIM13 classifier performance by behavior. Precision, recall, and F1 scores by behavior as a function of the proportion of data used to train the classifier on the CRIM13 dataset. Lines and shaded regions indicate mean and standard error, respectively, across 10 random splits of the data.

**Figure S4.**
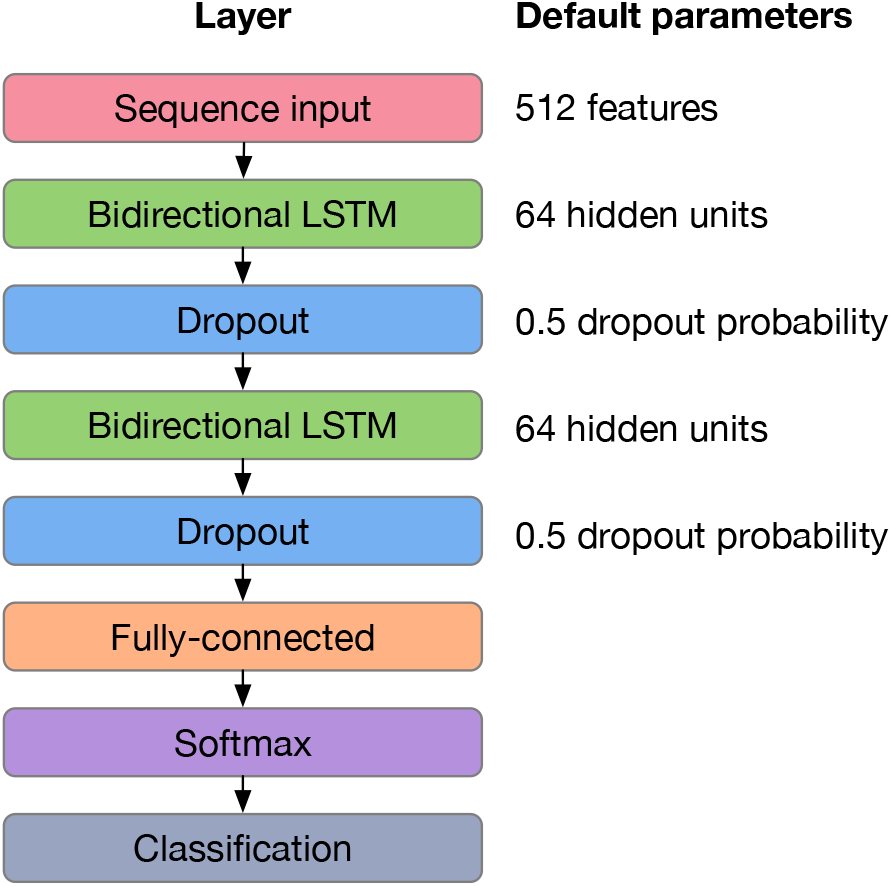
Schematic of the classification model. In our analysis, we use recurrent neural network that accepts input sequences with a dimensionality of 512. The sequence input layer is followed by a pair of bidirectional LSTM (64 hidden units) and dropout (with probability 0.50) layers. A fully connected layer accepts output from the second dropout layer and is followed by a softmax layer and a sequence-to-sequence classification layer, which returns the final output (i.e., a set of behavioral labels corresponding to each frame in the input).

